# SeSaMe PS Function: Functional Analysis of the Whole Metagenome Sequencing Data of the Arbuscular Mycorrhizal Fungi

**DOI:** 10.1101/2020.05.20.107235

**Authors:** Jee Eun Kang, Antonio Ciampi, Mohamed Hijri

## Abstract

In this article, we introduce a novel bioinformatics program- SeSaMe PS Function (<u>S</u>pore associated <u>S</u>ymbiotic <u>M</u>icrobes <u>P</u>osition <u>S</u>pecific Function) - for position-specific functional analysis of short sequences derived from metagenome sequencing data of the arbuscular mycorrhizal fungi. The unique advantage of the program lies in databases created based on genus-specific sequence properties derived from protein secondary structure, namely amino acid usages, codon usages, and codon contexts of three codon DNA 9-mers. SeSaMe PS Function searches a query sequence against reference sequence database, identifies three codon DNA 9-mers with structural roles, and dynamically creates the comparative dataset of 54 microbial genera based on their codon usage biases. The program applies correlation Principal Component Analysis in conjunction with K-means clustering method to the comparative dataset. Three codon DNA 9-mers clustered as sole member or with only a few members are often structurally and functionally distinctive sites that provide useful insights into important molecular interactions. The program provides a versatile means for studying functions of short sequences from metagenome sequencing and has a wide spectrum of applications.

## Introduction

Arbuscular mycorrhizal fungi (AMF) are plant root colonizing symbiotic microorganisms that promote plant growth and improve soil quality [1–3]. AMF increase the effectiveness of phytoremediation and improve crop yields in agroecosystems [1,4–10]. Despite the importance of AMF, their genetics is poorly understood, due in large part to their coenocytic multinucleate nature and strict symbiotic partnership with plants [11]. A number of studies reported strong evidence that AMF interact closely-tightly adhering to the surface or in the interior of mycelia and spores- or loosely with a myriad of microorganisms covering major bacterial and fungal taxa [6,12–16]. These microorganisms can be removed from AMF by using cocktails of antibiotics in axenic cultivation systems [17]. Yet, only few AMF taxa are able to be cured and cultivated *in vitro*, and most successful isolates in such systems mainly belong to the genus *Rhizoglomus* [18]. Given that the majority of AMF have not been successfully cultured axenically, it is possible that AMF may be meta-organisms, inseparable from their bacterial and fungal partners.

Whole genome sequencing (WGS) of AMF taxa has been achieved exclusively from those grown *in vitro*. Although they provide important insights into AMF genetics, they have limitations in serving as reference genome due to large intra and inter isolate genome variations [19,20]. Furthermore, sequence analysis of the WGS of AMF taxa grown *in vivo*, typically in a pot culture with a host plant, can be challenging because the sequencing data contain a large proportion of sequences belonging to AMF associated microorganisms; the WGS data of AMF represent a complex metagenome. However, they provide invaluable information about the associated microbial community because a great majority of the associated microorganisms cannot be clustered in laboratory conditions. Taxonomic classification of the whole metagenome sequencing (WMS) data is essential for studying AMF genomics and their interactions with the associated microorganisms. We introduced the bioinformatics program-SeSaMe (<u>S</u>pore associated <u>S</u>ymbiotic <u>M</u>icrobes) - for taxonomic classification of the WMS of AMF. In this article, we introduce a novel bioinformatics program- SeSaMe Position Specific Function (SeSaMe PS Function). It predicts important position-specific functional sites in a query sequence, based on amino acid usages, codon usages, and codon contexts of three codon DNA 9-mers derived from protein secondary structures extracted from Protein Data Bank (PDB) (rcsb.org) [22].

Recent studies have documented the multiple regulatory roles of codon usage and of codon context in transcription and translation (e.g., regulation of gene expression, diversification of gene products, translational efficiency and accuracy, and protein degradation efficiency) [23–30]. Several studies have emphasized the regulatory roles of codon usage and codon context of multiple consecutive codons [24,29,30]. In addition, synonymous codons are believed to be a key factor in determining the active folding state of a gene product in response to environmental changes. One recent study showed that a gene with multiple synonymous mutations produced a protein with increased tolerance to abiotic stresses [31]. Moreover, non-optimal codons serve specific roles in regulating circadian rhythms in response to changes of environmental conditions [32,33]. Therefore, codon usage and codon context must have been playing important roles in the adaptation of microorganisms to abiotic stresses [34,35]. We are beginning to scratch the surface of the regulatory roles of codon usage and codon context, and these studies appear to be just a tip of iceberg.

The main variable of the program-trimer usage bias- takes usages and contexts of both amino acids and nucleotides into consideration; it is the product of amino acid usage and three codon usage of a three codon DNA 9-mer. Generally, trimer usage bias has a broad range of variations among taxonomic groups but low variation among microorganisms belonging to the same taxonomic group. Trimer usage bias reflects the important attributes of multiple consecutive codons. Codon composition-i.e., codon context of three consecutive codons- is an important determinant of properties of mRNA structure that plays key roles in transcription and translation. Codon usage is associated with pauses in translation and determines biochemical properties of gene products. Both of the attributes affect protein folding.

SeSaMe PS Function identifies three codon DNA 9-mers with structural roles in a query sequence, and dynamically creates comparative dataset based on their trimer usage biases that are retrieved from 54 genus-specific bias databases (Figure 1). SeSaMe PS Function applies correlation Principal Component Analysis (PCA) in conjunction with K-means clustering method (PCA-Kmeans) to the comparative dataset. It enables users to identify three codon DNA 9-mers with distinctive characteristics: outliers. Outliers are often important position-specific functional sites that provide useful insights into molecular interactions.

**Figure 1.**
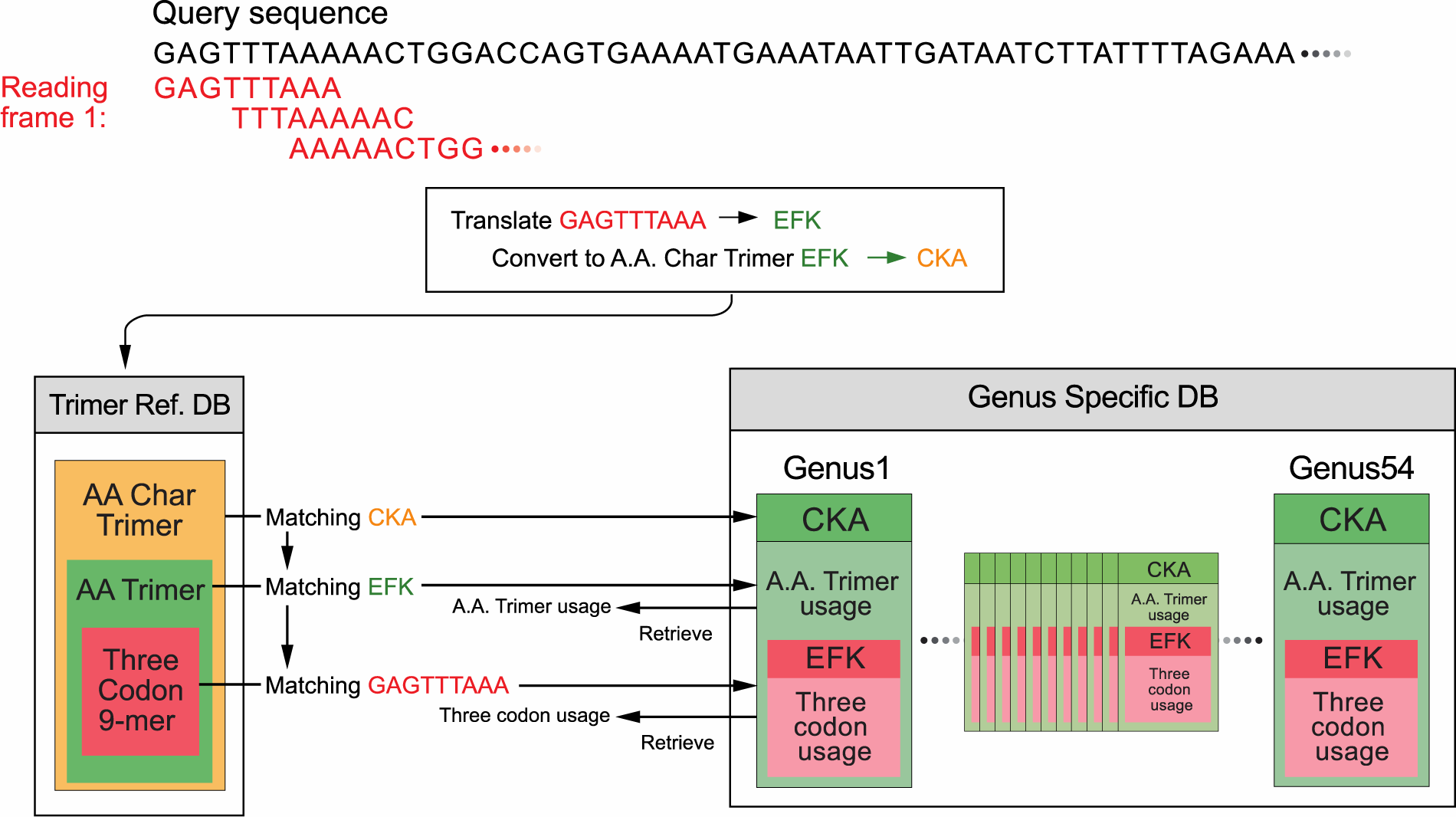
Dynamic creation of comparative dataset per query sequence. The program uses a query sequence to search matching A.A. Char Trimers, A.A. Trimers, and Three Codon DNA 9-mers in Trimer Ref. DB, and retrieves the A.A. Trimer usages of the matching A.A. Trimers and the three codon usages of the matching Three Codon DNA 9-mers from 54 Genus Specific DBs. It calculates the trimer usage biases of the matching Three Codon DNA 9-mers, and dynamically generates comparative dataset for the query sequence.

In this article, we analyzed one example sequence to demonstrate how to use the program for studying the structure and the function of a query sequence: one of the program’s various applications. The program helped to identify the outliers with potentially important functions such as ones involving in binding activities. Existing bioinformatics programs predicted that most of the outliers identified in the example sequence belonged to stem-loops, stems, and stem transitions in mRNA structures. Some of the outliers were matched to elements that play roles in promotor regions or in cis-regulatory mechanisms [36,40–42]. Other bioinformatics programs predicted that the example sequence may bind to DNA/RNA [22,39]. These results suggest that the outliers may contribute to binding activities in undiscovered mechanisms that may have attributes similar to cis-regulatory mechanism.

A majority of existing bioinformatics tools for position-specific sequence annotation rely on sequence alignments, which have low sensitivity toward hypervariable sequence motifs with flexible structures and various functions. Although they provide important information about a query sequence, their usage is limited to a particular set of motifs with known functions. In contrast, as SeSaMe PS Function employs PCA to identify outliers based on internal structure of comparative dataset that are created solely from a query sequence, it may reveal important molecular interaction sites not only in known but also in undiscovered mechanisms. It has been only several decades since advances have been made in molecular biology. Therefore, it is believed that only a small fraction of mechanisms in biological system have been discovered. SeSaMe PS Function provides useful tools for studying functions of short sequences from metagenome sequencing data. It is available for download free of charge at www.journal.com and at www.fungalsesame.org.

## Methods

### Database design and comparative dataset creation

The databases were originally created for the metagenome taxonomic classifier-SeSaMe, and then incorporated into SeSaMe PS Function. While NCBI offered a large number of completely sequenced bacterial genomes, only a small number of fungal genomes were completely sequenced. The completely sequenced genomes of 444 bacteria and of 11 fungi, known to be present in soil, were downloaded and assigned into 45 bacterial and 9 fungal genera, respectively. CDS database per genus was created based on CDS list provided by NCBI, JGI, or Tisserant et al. [19].

The program consists of two types of databases and a PCA-Kmeans method. 7,674 amino acid trimers, found in protein secondary structures in PDB, were selected and then assigned to the sequence variable-A.A. Trimer in the trimer reference sequence database (Trimer Ref. DB) (Figure 2). Amino acid characteristic (A.A. Char) is defined as a group of amino acid(s) with similar property(s), and consists of 12 groups: A (Lysine (K), Arginine (R)), B (Histidine (H)), C (Aspartic acid (D), Glutamic acid (E)), D (Serine (S), Threonine (T)), E (Asparagine (N), Glutamine (Q)), F (Cysteine (C)), G (Glycine (G)), H (Proline (P)), I (Methionine (M)), J (Alanine (A), Isoleucine (I), Leucine (L), Valine (V)), K (Phenylalanine (F), Tryptophan (W), Tyrosine (Y)), and L (stop codons). Trimer Ref. DB consists of three sequence variables that form a three level hierarchy: amino acid characteristic trimer (A.A. Char Trimer), A.A. Trimer, and Three Codon DNA 9-mer (Figure 1).

**Figure 2.**
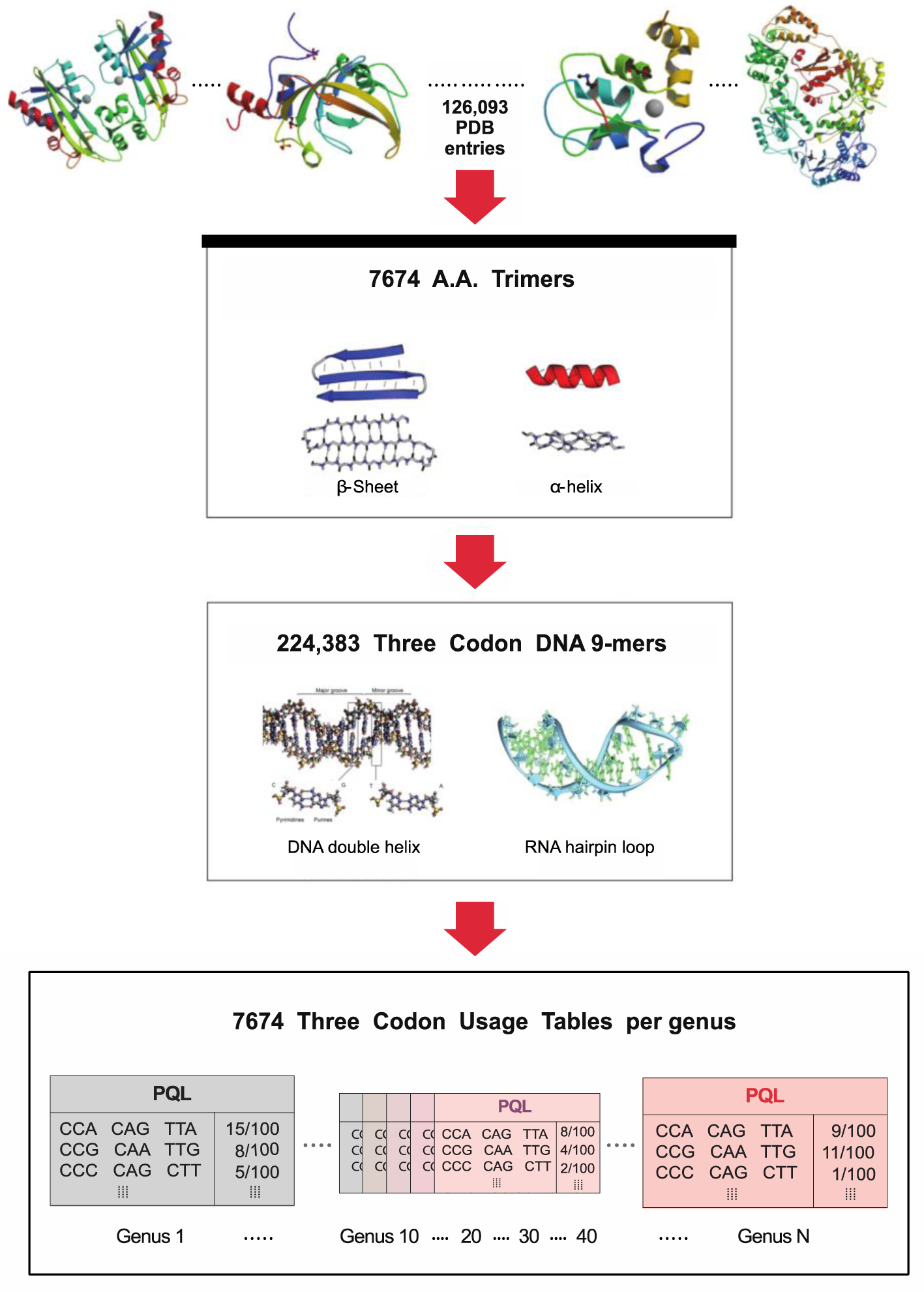
Database Design. A large number of PDB entry files were processed to extract 7674 A.A. Trimers-subunits of protein secondary structures. A table of three codon usage was created per A.A. Trimer and per genus in Genus Specific DB. *Note*: The PDB IDs of the protein structures in the top of the figure from left are 2ZTI, 3DWH, 2VSL, and 3DRP. Their citations are included in the reference section. Image of the protein secondary structures and the nucleotide structures in the first box and in the second box are from https://en.wikipedia.org/wiki/Alpha_helix and https://en.wikipedia.org/wiki/RNA, respectively.

Genus-specific usage bias database (Genus Specific DB) contains the numerical variables-A.A. Trimer usage of A.A. Trimer and three codon usage of Three Codon DNA 9-mer. The main numerical variable, trimer usage bias, is calculated by multiplying A.A. Trimer usage by three codon usage. There are 54 Genus Specific DBs where each Genus Specific DB consists of 1,296 A.A. Trimer Usage Tables and 7,674 Three Codon Usage Tables created based on the CDS database (Figure 2).

For each reading frame of a query sequence, the program uses a query sequence to search against Trimer Ref. DB, identifying matching A.A. Char Trimers, A.A. Trimers, and Three Codon DNA 9-mers. It retrieves the trimer usage biases of the matching Three Codon DNA 9-mers from 54 Genus Specific DBs, and creates a comparative dataset of 54 genera (Figure 1). The input matrix to the correlation PCA method is the comparative dataset with 54 genera in rows (observations) and the matching Three Codon DNA 9-mers in columns. The input matrix will be called hereafter Z (I x J).

### Annotation for catalytic and allosteric sites

According to Catalytic Site Atlas and Allosteric Database, A.A. Trimers were divided into 4 subgroups based on the property of their second amino acid-catalytic site (CSA), allosteric site (ASD), both CSA and ASD (Both), and none of them (None) [47,48]. An A.A. Trimer in CSA, ASD, or Both groups was annotated with the list of functions of PDB molecules that contained the A.A. Trimer. This feature is for making inferences about functionality not of a query sequence but of its A.A. Trimers.

### Implementation of the correlation PCA-Kmeans method

The correlation PCA method was implemented based on the method reported by Abdi et al. (2010) [49], which provides important definitions and multiple examples to help readers understand the concepts underlying PCA [49]. Interpretation of the result from SeSaMe PS Function also relies on Abdi et al. (2010) because eigenvalue decomposition is mathematically closely related to singular value decomposition and has similar underlying concepts. Pearson’s correlation method is applied to the centered Z (input matrix) and produces a correlation matrix X (J x J). Eigenvalue decomposition is applied to X and produces components. V is an eigenvector matrix with J x J dimensions and is also called a loading matrix.

#### Loadings: elements of the loading eigenvector matrix V

The program calculates an eigenvector matrix V. Loading is defined as the element of V. V has matching Three Codon DNA 9-mers in rows and the same number of components in columns. The program examines loadings on components whose sum accounts for 80% of inertia (80% components) in addition to loadings on the first principal component and the second component (the First/Second components) [49]. The program creates two different input matrices based on V called L1 and L2. They have the same number of Three Codon DNA 9-mers in rows. L1 has 80% components in columns while L2 has the First/Second components in columns. The program separately applies the K-means clustering method (default k = 13) to L1 and L2.

#### Taxon scores of 54 genera in component spaces

The program calculates taxon scores of 54 genera observations. Taxon score matrix (I x J) results from multiplying centered Z by V. Inertia of a component is defined as a sum of squared taxon scores in corresponding component column [49]. The program creates two matrices based on taxon score matrix called T1 and T2. They have 54 genera observations in rows. T1 has 80% components in columns, while T2 has the First/Second components in columns. The program separately applies the K-means clustering method (default k = 10) to T1 and T2.

### Program availability

The program was implemented in Java programming language (www.java.net, www.oracle.com (Java8)). We used the Pearson’s correlation, the eigenvalue decomposition, and the K-means clustering in the Apache Commons Math3 library (3.3). The program requires the Apache Commons Math3 (3.3) and IO (2.4) libraries (www.apache.org). The program has been made to run on Linux/ Unix operating systems, packaged into an executable Java JAR Jfile, and tested and confirmed to work on Linux system-CentOS Linux 7 (www.centos.org). The program that is being introduced in this article is version 1 and was implemented with the correlation PCA only. The program (version 2) was implemented both with the covariance PCA and with the correlation PCA. They have been used at the Biodiversity Center, Institut de Recherche en Biologie Végétale, Département de Sciences Biologiques, Université de Montréal. They are available for download free of charge at www.journal.com and www.fungalsesame.org. There are no restrictions for using the programs by academic or non-academic organizations as long as a user complies with the license agreement.

### Input, output, and options

The program has a command-line interface. Input files should contain DNA sequence(s) in fasta format. It requires a command-line argument-input file path. SeSaMe PS Function produces three different types of outputs per query sequence. One is the standard PCA output: the sequence information of matching Three Codon DNA 9-mers, the percentage of an explained inertia by a component, and the contribution of an observation to a component [49]. Another is the loading cluster output with the loading information. Each Three Codon DNA 9-mer is annotated with subgroups-CSA/ ASD/ Both/ None and the functions of PDB molecules. The other is the genus cluster output with the taxon scores. It should be noted that the cluster result is different for every run, because the K-means clustering method in the Apache Commons Math library randomly chooses initial centers for multiple iterations to decrease chances of poor clustering.

SeSaMe PS Function version 1 and version 2 have an option to specify the k parameter in the K-means clustering method both for genus clusters and for loading clusters (e.g., 11_15). The program version 2 has an additional option called “auto”. If a user wants to run SeSaMe PS Function for a large number of query sequences with varying lengths, he can use the prefix “auto” to set the k parameter for loading clusters according to a simple equation: the number of matching Three Codon DNA 9-mers divided by a user specified number. For example, if a user gives the following option “auto_14_8”, it will automatically set one eighth of the number of matching Three Codon DNA 9-mers as the k parameter for loading clusters while it will set 14 as the k parameter for genus clusters. A suitable k value may vary widely depending on the length and the complexity of a query sequence. User can supply the option following the input file path (e.g., /home/input-file auto_14_8).

### Demonstration of the program usage

#### Selection of the example sequence

We selected 25 correctly predicted sequences out of 100 AMF CDS test sequences that were used for evaluating the accuracy of the metagenome taxonomic classifier, SeSaMe. From 25 sequences, we selected one example sequence that had the largest number of Three Codon DNA 9-mers where AMF had the highest trimer usage bias among 54 genera. The example sequence is TGAGTTTAAAAACTGGACCAGTGAAAATGAAATAATTGATAATCTTATTTTAGAAATGCAATTAAAAATTAATAGTACATATGATAAAATAGTTGAATGGATACCATACAATCAGTTTATTAACATTAACGAAATAGGAAAAGTTGGTGATAATACTGCTGTATATTCAGCAATATGGAAAAATGGTCCACTATATTATAGAAAGAAATGGATAAGGAAATCCAATGAAAAAGTTGTATTAAATTACTTAACATTAGATATTAAGGAATT.

#### Outlier’s unique pattern of the trimer usage bias and of the three codon usage

Landscape pattern is the comparison of 54 genera based either on the trimer usage bias or on the three codon usage of a Three Codon DNA 9-mer. It provides an accurate way to estimate the relative measure of the usage information across 54 genera. In this article, we abbreviate Three Codon DNA 9-mer according to the order of its position in DNA sequence and its A.A. Trimer (Table S1). For example, AATACTGCT is the 51st matching Three Codon DNA 9-mer and encodes for the amino acids NTA. Because the program is zero-based, its abbreviation is 50 NTA. Graphs showing the landscape patterns of three codon usages and of trimer usage biases retrieved from 54 genera were generated for 17 EMQ and 67 KKW and for 18 MQL and 3 NWT, respectively.

#### Comparison of the frequencies of a nucleotide among 13 loading clusters

We counted the frequencies of the nucleotide-adenine (A) in each of the individual Three Codon DNA 9-mers and applied a one-way ANOVA test to compare the means among 13 clusters. We repeated the same process for the nucleotides cytosine (C), guanine (G), and thymine (T).

#### Comparison between the trimer usage bias and the three codon usage in functional segment

We assigned matching Three Codon DNA 9-mers into functional segments (FSs) based on the loading clusters with 80% components and based on the prediction result of the protein secondary structure from a bioinformatics tool-SCRATCH [50].

We created two matrices per FS; one was based on the three codon usage, and the other was based on the trimer usage bias. Each matrix consisted of the usage information of the matching Three Codon DNA 9-mers retrieved from 54 genera; it had the Three Codon DNA 9-mers of an FS in rows and the 54 genera in columns. After centering the matrix, we applied Pearson’s correlation to each matrix to yield a correlation matrix (I x I), and calculated the mean of the correlations per pair of taxonomic groups-Clostridia, Bacilli, Oscillatoriophycideae, Nostocales, Acidobacteria, Alphaproteobacteria, Betaproteobacteria, Deltaproteobacteria, Gammaproteobacteria, AMF, Agaricomycotina, and Pezizomycotina. From the mean of the correlations of a pair of genera belonging to the same taxonomic group in each FS, we calculated the mean and the standard deviation per taxonomic group. In the same way, we calculated the mean of the correlations for pairs of taxonomic groups-Firmicutes, Cyanobacteria, Proteobacteria, Actinobacteria, AMF, a group of 7 Dikarya, and *Phanerochaete* in each FS.

### Results of the selected analysis

#### Loading clusters

The example sequence had 270 bp. When we ran the metagenome taxonomic classifier-SeSaMe- with the example sequence, it had the highest trimer usage probability score in the 2nd reading frame translation. It had 87 matching Three Codon DNA 9-mers in the 2nd reading frame translation. The PCA method applied to the comparative dataset showed that 51 components represented 80% components, while the First/Second components explained approximately 29% of total inertia.

The K-means clustering method (k = 13) applied to the loadings of 80% components identified outliers: 14 Three Codon DNA 9-mers in 12 clusters. Ten clusters had a sole member (50 NTA, 63 LYY, 72 KSN, 4 WTS, 69 WIR, 73 SNE, 24 STY, 30 VEW, 80 NYL, and 51 TAV) while two clusters had two members (33 IPY and 61 GPL and 39 INI and 86 IKE). One major cluster had 73 members.

Structural homology search in PDB and inference of DNA-binding residues in DRNApred suggested that the example sequence may be a DNA/RNA binding protein [22,39]. We used the outliers to search publicly available bioinformatics databases containing DNA motifs with known functions. RSAT indicated that the outlier and its adjacent Three Codon DNA 9-mer (4 WTS and 3 NWT) were matched to motifs involved in cis-regulatory mechanisms, one in the + strand and the other in the – strand [40]. BPROM (Prediction of bacterial promoters) predicted that the outliers 30 VEW and 33 IPY were promoter-related elements [41]. GPMiner indicated that three outliers (4 WTS, 33 IPY, and 61 GPL) were matched to statistically significant over-represented oligonucleotides in the promoter region [42]. RNA structure prediction tools predicted that most outliers formed stem-loops, stems, and transition routes to stem in mRNA structure of the example sequence (Figure S1) [36]. A large number of studies have documented stem-loop and stem structures in mRNAs as important regulatory sites and binding sites [37,38]. Considering that we are just beginning to understand the regulatory roles of codon usage and codon context, considerable portions of outliers and their adjacent Three Codon DNA 9-mers identified by the program may serve important roles in undiscovered mechanisms.

The loading clusters with the First/Second components based on the trimer usage bias are shown in Table S1. It should be noted that Table S1 indicates the three codon usages for comparison purpose, which will be discussed in another section. The loadings of Three Codon DNA 9-mers with the catalytic or with the allosteric site in the second amino acid were plotted on the space of the First/Second components (Figure 3). A majority of Three Codon DNA 9-mers where Firmicutes, Cyanobacteria, Rickettsia, and AMF had the highest three codon usage were aggregately located on the far-right side (Figure 3). In contrast, those where Deltaproteobacteria, Gammaproteobacteria, and Actinobacteria had the highest three codon usage were dispersed across the left side and the middle of the graph. For example, 3 NWT where Kocuria had the highest value was located on the far-left side (Table S1).

**Figure 3.**
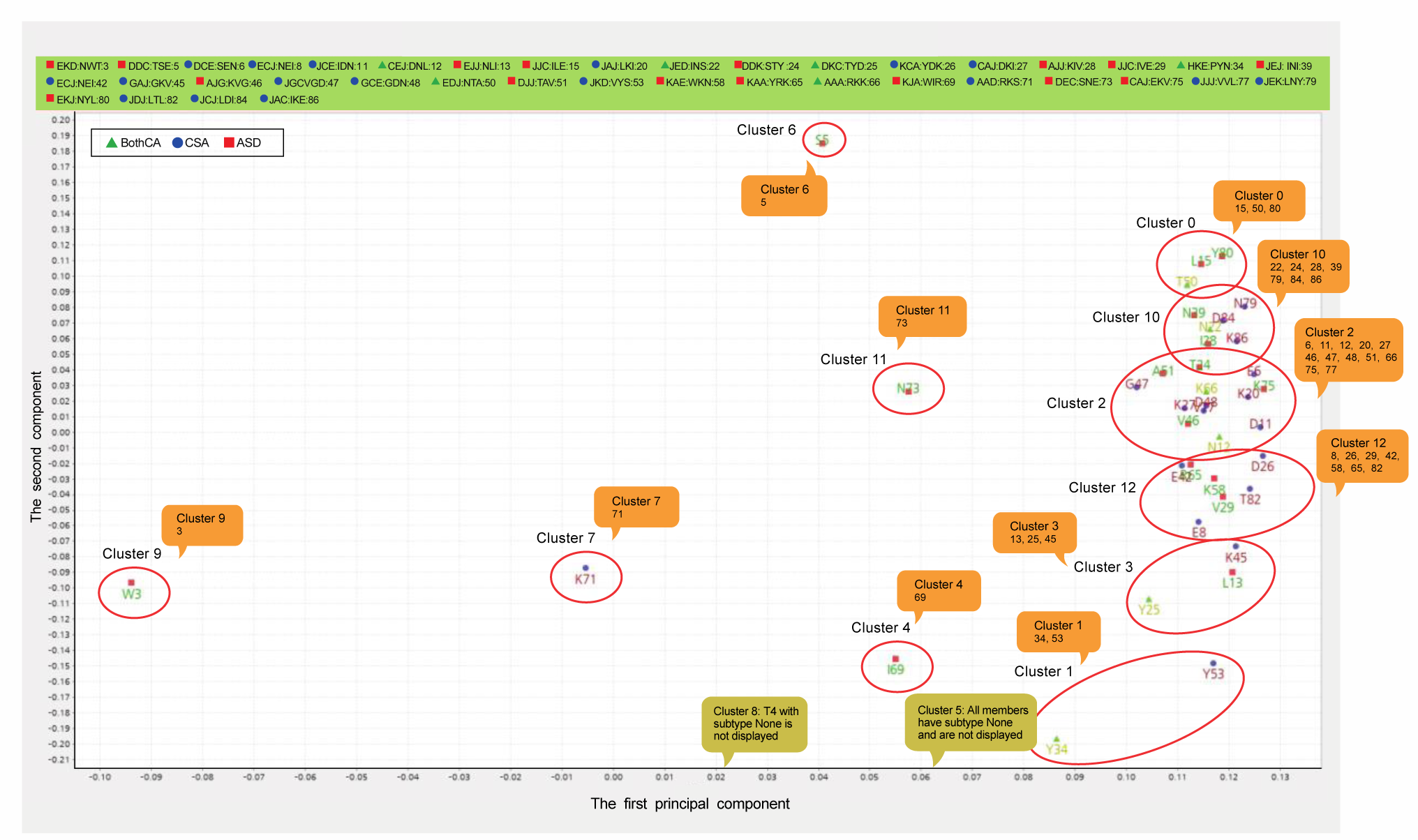
Loading clusters of the example sequence. The figure shows elements of the loading matrix V on the space of the First/Second components. Note: Abbreviation: a name of Three Codon DNA 9-mer was abbreviated to the second amino acid of its A.A. Trimer. For example, Three Codon DNA 9-mer (AACTGGACC), encoding for the A.A. Trimer NWT, was abbreviated to W (Table S1). A digit next to the abbreviation indicates the order of its position in the example sequence. A digit in the colored box is abbreviation of Three Codon DNA 9-mer. For example, 22 under Cluster 10 in the box stands for ATTAATAGT that encodes for the A.A. Trimer INS and whose order of the position is 22. CSA, ASD, Both, and None stands for catalytic site, allosteric site, both catalytic and allosteric site, and none of these sites, respectively.

#### Genus clusters

The genus clusters based on 80% components indicated that genera with close phylogenetic relationships were assigned to the same cluster. In the scatter plot of taxon scorers on the space of First/Second components, Firmicutes, Cyanobacteria, *Rickettsia*, and AMF that frequently had higher trimer usage biases were located on the right while most members of Actinobacteria and Proteobacteria (cluster 1) that frequently had lower values were located on the far-left side (**Figure 4**).

**Figure 4.**
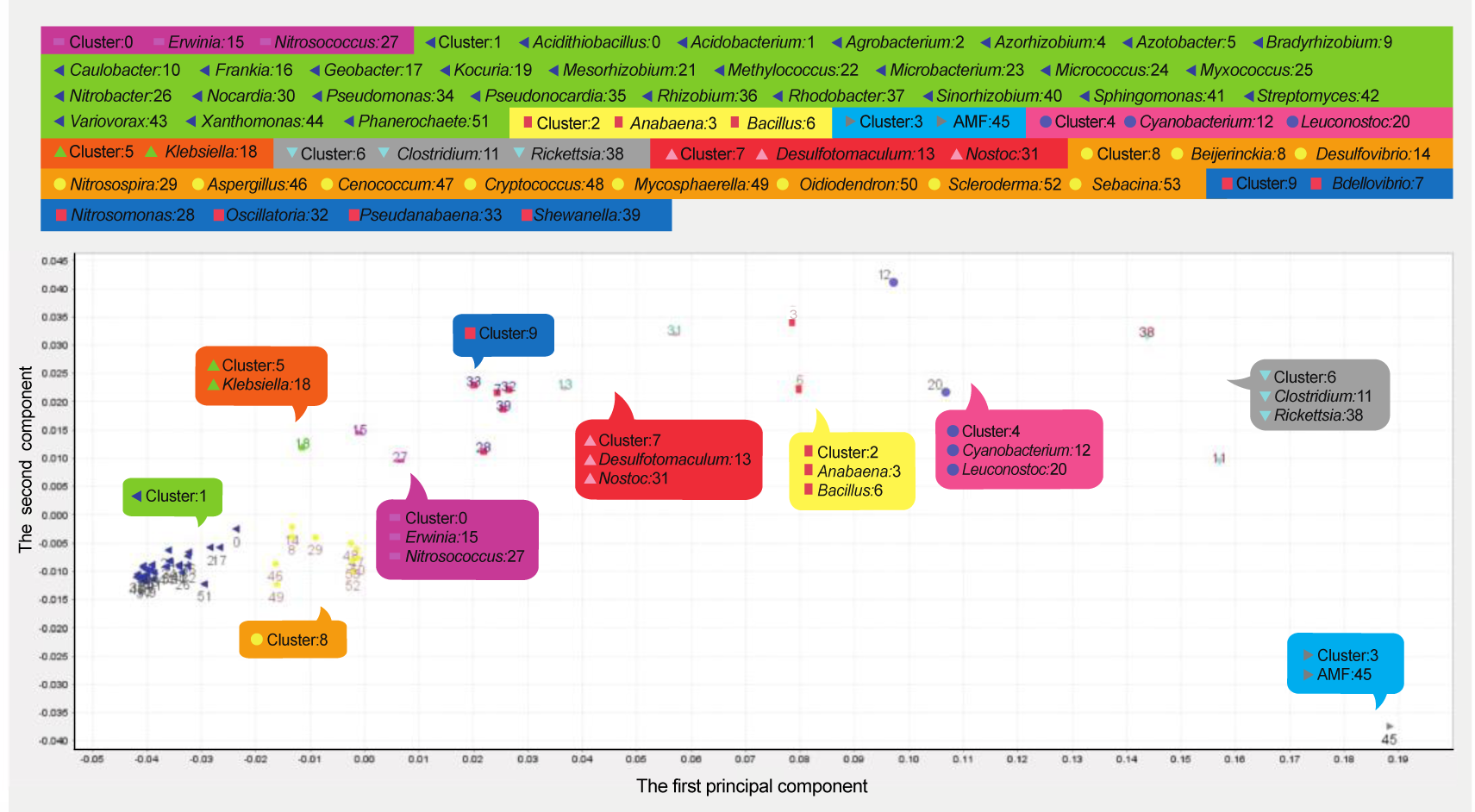
Genus clusters of the example sequence. Taxon scores of 54 genera are plotted on the space of the First/Second components.

#### Outlier’s unique landscape pattern of trimer usage bias and of three codon usage

For each of the Three Codon DNA 9-mers in loading clusters with the First/Second components, we ranked 54 genera in order of decreasing three codon usage. We then, ranked the Three Codon DNA 9-mers in each subgroup (CSA/ ASD/ Both/ None) of the clusters based on a maximum of the three codon usages (Table S1). The mean of the maxima was 0.256. AMF, Clostridium, and Rickettsia frequently had the maximum.

Most of Three Codon DNA 9-mers in the major cluster demonstrated similar landscape patterns of the three codon usage and of the trimer usage bias. For example, 17-EMQ-GAAATGCAA and 18-MQL-ATGCAATTA had the frequently demonstrated landscape pattern (Figures S2 and S3). Outliers had a unique landscape pattern; genera belonging to Dikarya had a higher value than AMF both in 67-KKW-AAGAAATGG and in 3-NWT-AACTGGACC (Figures 5 and S4).

**Figure 5.**
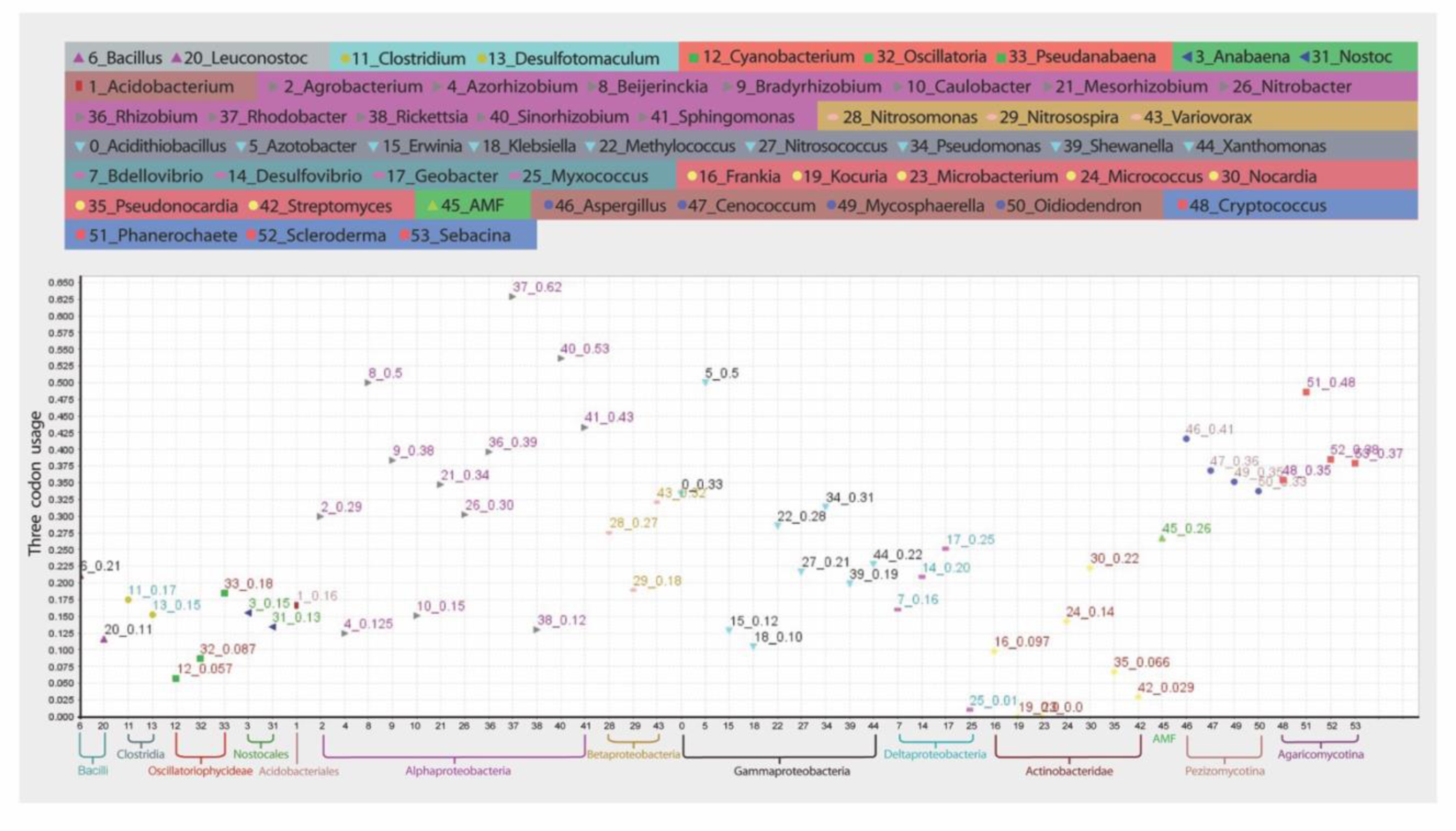
Landscape pattern of the three codon usage of 67-AAK-KKW-AAGAAATGG. 54 genera are arranged into 13 taxonomic groups.

#### Comparison of the frequencies of a nucleotide among 13 loading clusters

One-way ANOVA tests showed that the means of the frequencies of G and C in each of the individual Three Codon DNA 9-mers were significantly different among 13 clusters; F-statistics and p-value of A, T, G, and C among 13 clusters were 0.69 (0.76), 1.26 (0.26), 1.91 (0.047), and 3.09 (0.0014), respectively.

#### Comparison between the trimer usage bias and the three codon usage in FS

We merged some of the outliers in 12 clusters according to the proximity of their primary sequences, which resulted in 8 groups. The merged outliers were 50 NTA with 51 TAV, 33 IPY and 61 GPL with 63 LYY, and 69 WIR with 72 KSN and 73 SNE. This was done to simplify the analysis, and is not recommended for real case analyses. Examining the protein tertiary structure predicted by SCRATCH, we added another group (20 LKI), a member of alpha helix, which resulted in a total of 9 groups [50]. We assigned 87 Three Codon DNA 9-mers into 9 FSs according to the outliers: FS1: 4 WTS (from Three Codon DNA 9-mer 0 – 12); FS2: 20 LKI (alpha helix1: 13 – 21); FS3: 24 STY (22 – 29); FS4: 30 VEW (30 – 32); FS5: 33 IPY, 61 GPL, 63 LYY (33 – 35, 52 – 65); FS6: 39 INI, 86 IKE (36 – 41, 82 – 86); FS7: 50 NTA, 51 TAV (42 – 51); FS8: 69 WIR, 72 KSN, 73 SNE (66 – 73); FS9: 80 NYL (alpha helix2: 74 – 81) (Figure S5).

Generally, the mean of the correlations of a pair of genera belonging to the same taxonomic group was the highest in each taxonomic group for all 9 FSs (Tables S2 and S3). Table S4 shows the mean and the standard deviation of 9 FSs calculated from the mean of the correlations of a pair of genera belonging to the same taxonomic group in a FS.

The mean of the correlations of a pair of taxonomic groups based on three codon usage (left) and the mean based on trimer usage bias (right) are shown in Table S3. Most of them had strong correlations in both alpha helices – FS2 and FS9. This may suggest that roles of amino acids and of codons in alpha helices may be relatively more conserved across taxonomic groups due to functional and structural constraints compared to those in random coils and loops of which flexible structures are equipped for a variety of functions.

#### Comparable properties of 25 selected sequences in AMF CDS test set

In order to show that the program provided outliers of Three Codon DNA 9-mers in loading clusters (80% components) not only in the example sequence but also in all 25 sequences, we included the cluster results of 5 additional query sequences. Genus clusters and loading clusters of the sequences are shown in Supplementary Tables 5 and 6, respectively. The early diverged bacteria and AMF were often clustered as sole members or with each other. A great majority of Three Codon DNA 9-mers were grouped together into one major cluster, while outliers were clustered as sole members or with only one other member.

## Future work

Both the trimer usage bias and the three codon usage showed comparable landscape patterns. It may suggest that they provide a coherent explanation for as yet undiscovered regulatory mechanisms where codons play key roles in folding mRNAs and proteins. Recent studies have documented that codon usage and mRNA structure regulate protein folding [24,25,27,30]. For example, some studies showed association between rare codons or double stranded mRNA structures and a decrease of translational speed [25,30]. Other studies have documented relationships between protein secondary structure and mRNA structure; double stranded mRNA regions tend to encode alpha helix and beta-strand while single stranded mRNA regions tend to encode random coils [51,52]. However, the roles of the codons involved in these rules may vary widely across taxonomic groups. Nevertheless, while we need solved structures of proteins and RNAs across various taxa, they are mostly from a small number of model organisms.

Therefore, it is challenging to connect mRNA structures with their corresponding protein structures in metagenome sequencing data. We may be able to improve SeSaMe PS Function by incorporating mRNA structure information with an algorithm that dynamically predicts single and double stranded regions in a query sequence. This may add structurally distinctive properties of Three Codon DNA 9-mers across taxonomic groups, which may increase the capacity of taxonomic classification and at the same time, provide important information on the structure and the function of a query sequence.

## Authors’ contributions

KJE designed the program and implemented it using Java programming language. CA gave advice on what needs to be included in result of the correlation PCA-Kmeans method in terms of statistics, and helped to draft the manuscript. HM gave advice on developing the method, and helped to draft the manuscript. All authors read and approved the final manuscript.

## Competing interests

The authors have declared no competing interests.

## Acknowledgments

The authors gratefully acknowledge AFE (Éducation et de l’Enseignement supérieur Quebec), FESP (Faculté des études supérieures et postdoctorales de l’UdeM), and IRBV (Institut de Recherche en Biologie Végétale de l’Université de Montréal) for awarding scholarships to KJE. We thank David Morse for editing and commenting on the manuscript.

## Figure Legends

**Supplementary Figure S1 Outliers in the predicted mRNA secondary structures**

The structures were generated by the bioinformatics program-RNAstructure (https://rna.urmc.rochester.edu/RNAstructureWeb/Servers/Predict1/Predict1.html).

**A.** The example DNA sequence with T replaced with U was submitted as RNA. The secondary structure was predicted based on the algorithm called Fold. **B.** The example DNA sequence was submitted as DNA. The secondary structure was predicted based on the algorithm called MaxExpect.

**Supplementary Figure S2 Landscape pattern of the three codon usage of 17-CIE-EMQ-GAAATGCAA**

54 genera are arranged into 13 taxonomic groups.

**Supplementary Figure S3 Landscape pattern of the trimer usage bias of 18-IEJ-MQL-ATGCAATTA**

54 genera are arranged into 13 taxonomic groups.

**Supplementary Figure S4 Landscape pattern of the trimer usage bias of 3-EKD-NWT-AACTGGACC**

54 genera are arranged into 13 taxonomic groups.

**Supplementary Figure S5 FSs of the predicted protein tertiary structure**

The structure was predicted by the bioinformatics program-SCRATCH (http://scratch.proteomics.ics.uci.edu/). The PDB file format was converted to Cn3D format by another bioinformatics program-Vast (https://www.ncbi.nlm.nih.gov/Structure/VAST/vastsearch.html). The 3-dimensional structure was viewed by Cn3D (https://www.ncbi.nlm.nih.gov/Structure/CN3D/cn3d.shtml).

**Supplementary Table S1 Loading clusters according to the First/Second components**

A maximum of three codon usages among 54 genera and a genus with the maximum for each Three Codon DNA 9-mer is indicated in the column-three_max and three_genus, respectively.

**Supplementary Table S2 Correlations of a pair of genera based on trimer usage bias**

FS 1

FS 2

FS 3

FS 4

FS 5

FS 6

FS 7

FS 8

FS 9

**Supplementary Table S3 The mean of the correlations in 9 FSs**

The mean and the standard deviation of the correlations of a pair of genera were calculated based on three codon usage (left) and on trimer usage bias (right) in 9 FSs in taxonomic groups-bacterial phyla, AMF, a group of 7 Dikarya, and *Phanerochaete.*

**Supplementary Table S4 The mean and the standard deviation of the correlations of 9 FSs**

Based on three codon usage

Based on trimer usage bias

**Supplementary Table S5 Genus clusters of five additional sequences**

**Supplementary Table S6 Loading clusters of five additional sequences**

## Notes

http://www.fungalsesame.org/

