## Supplementary Figures S1-S5 for "SeSaMe PS Function: Functional Analysis of the Whole Metagenome Sequencing Data of the Arbuscular Mycorrhizal Fungi"

**Supplementary Information**
**Kang et al. Genomics, Proteomics $ Bioinformatics (Ref, GPB-D-17-00137)**

**Figure S1**

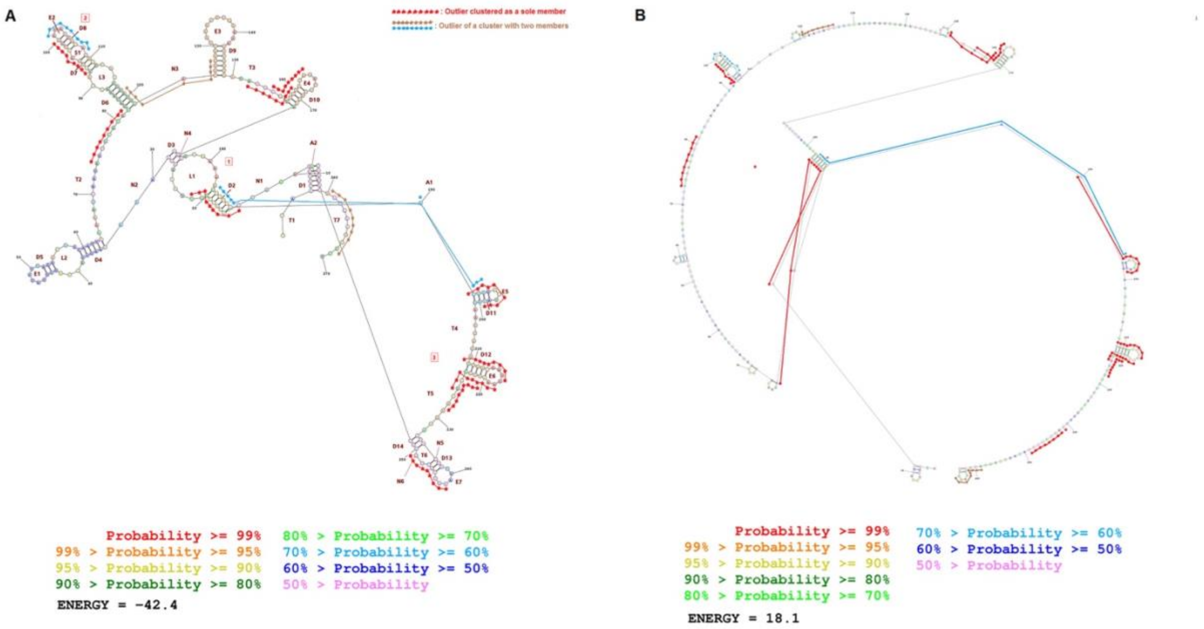


**Supplementary Figure S1 Outliers in the predicted mRNA secondary structures**
The structures were generated by the bioinformatics program- RNAstructure (https://rna.urmc.rochester.edu/RNAstructureWeb/Servers/Predict1/Predict1.html).
**A.** The example DNA sequence with T replaced with U was submitted as RNA. The secondary structure was predicted based on the algorithm called Fold. **B.** The example DNA sequence was submitted as DNA. The secondary structure was predicted based on the algorithm called MaxExpect.

**Figure S2**

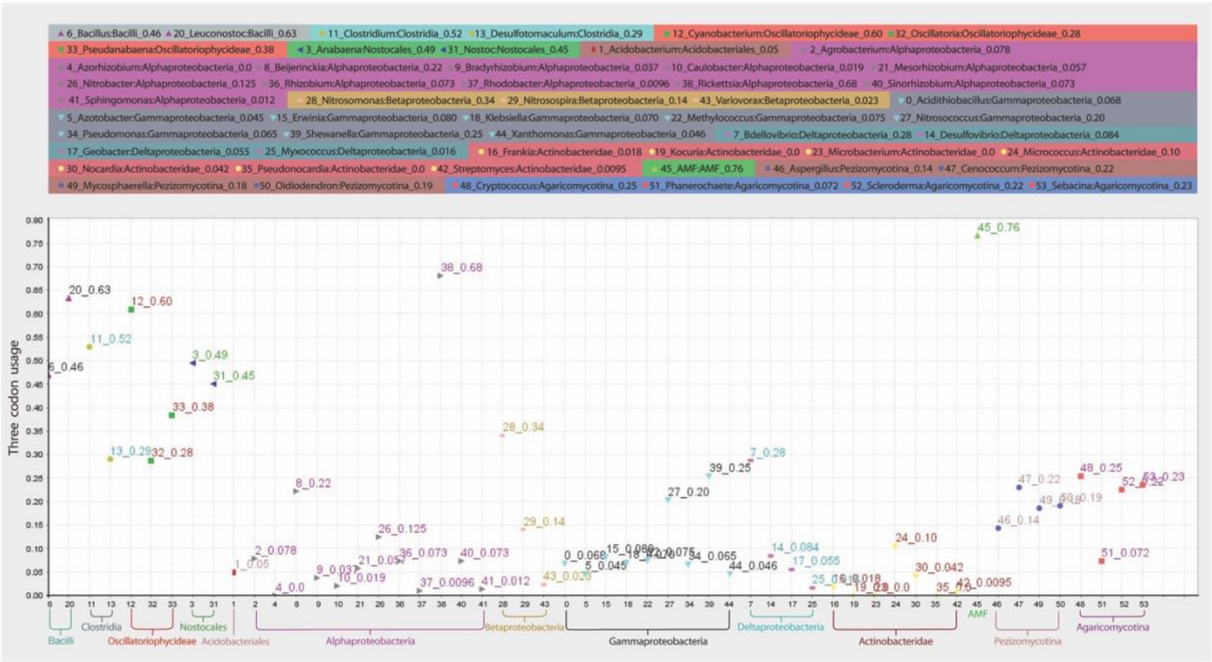


**Supplementary Figure S2 Landscape pattern of the three codon usage of 17-CIE-EMQ-GAAATGCAA**

54 genera are arranged into 13 taxonomic groups.

**Figure S3**

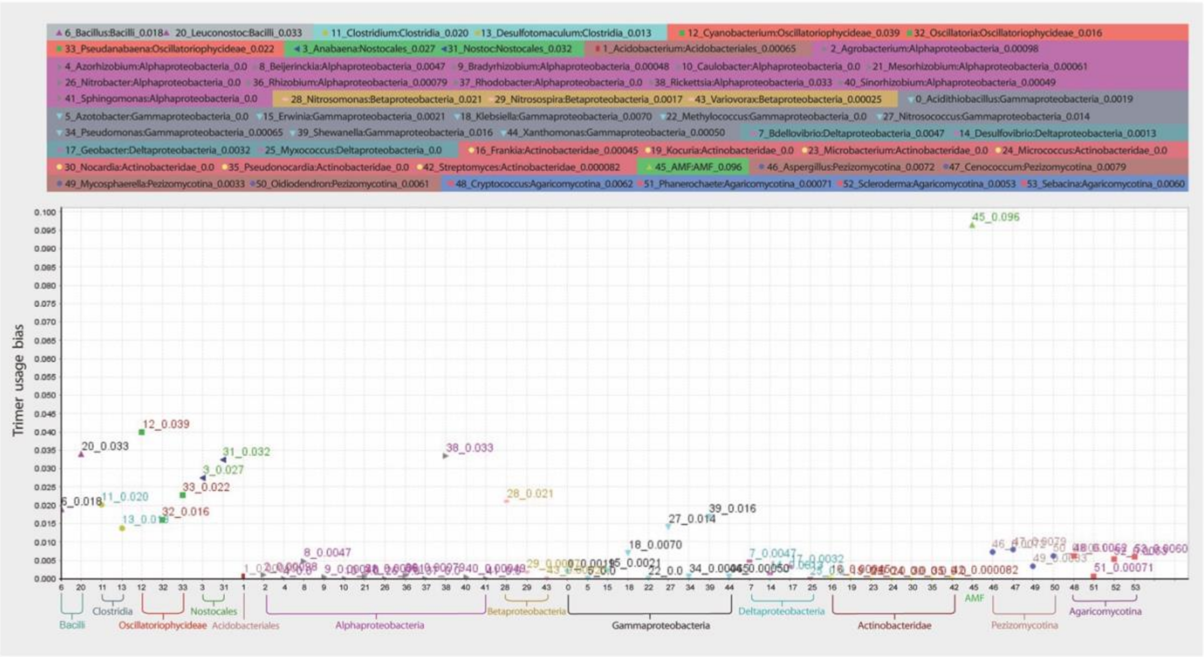


**Supplementary Figure S3 Landscape pattern of the trimer usage bias of 18-IEJ-MQL-ATGCAATTA**

54 genera are arranged into 13 taxonomic groups.

**Figure S 4**

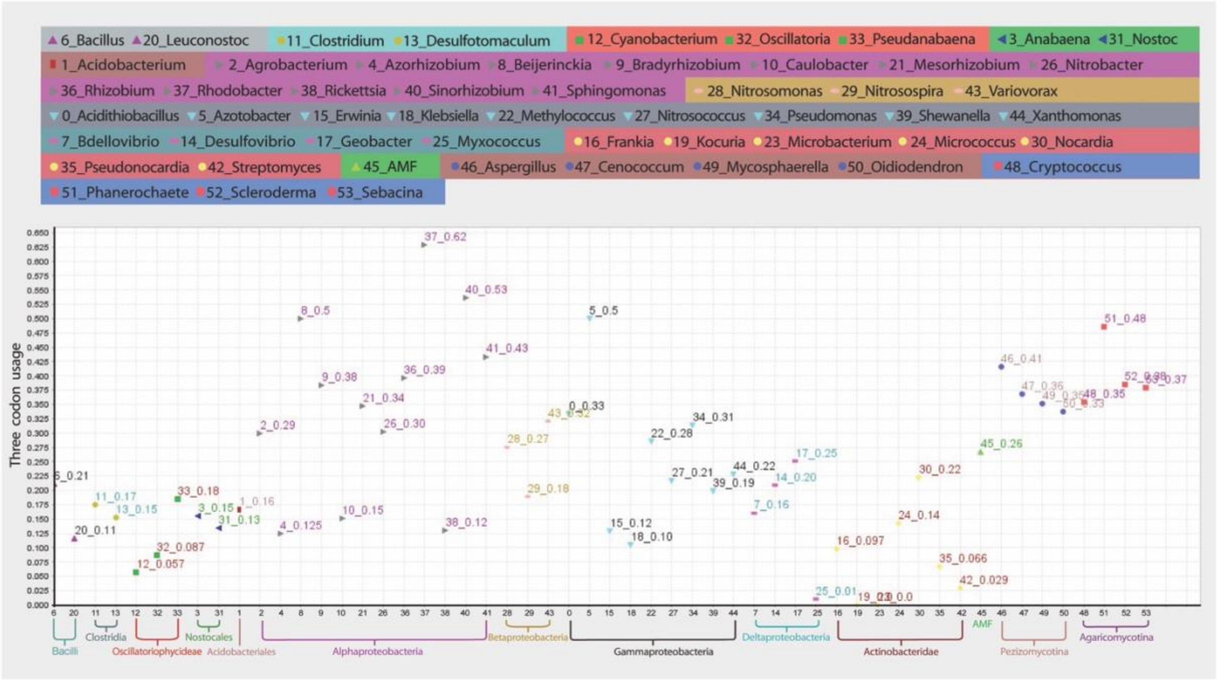


**Supplementary Figure S4 Landscape pattern of the trimer usage bias of 3-EKD-NWT-AACTGGACC**

54 genera are arranged into 13 taxonomic groups.

**Figure S5**

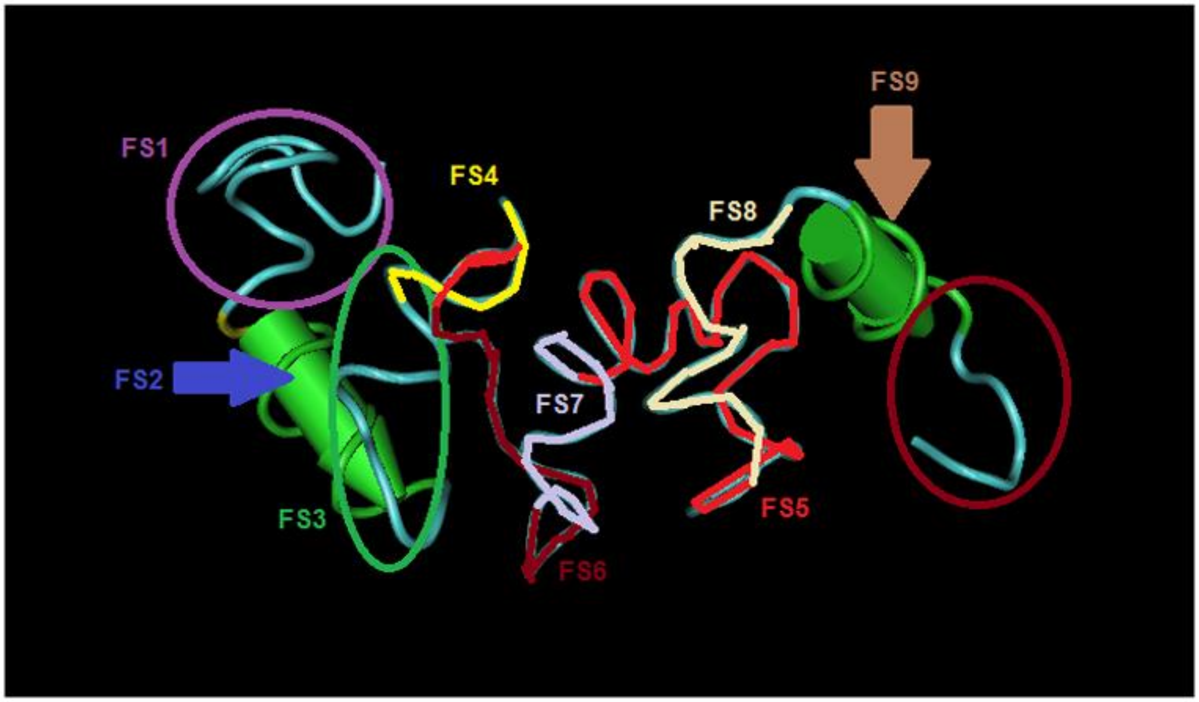


**Supplementary Figure S5 FSs of the predicted protein tertiary structure**

The structure was predicted by the bioinformatics program- SCRATCH (http://scratch.proteomics.ics.uci.edu/). The PDB file format was converted to Cn3D format by another bioinformatics program- Vast (https://www.ncbi.nlm.nih.gov/Structure/VAST/vastsearch.html). The 3-dimensional structure was viewed by Cn3D (https://www.ncbi.nlm.nih.gov/Structure/CN3D/cn3d.shtml).

**Supplementary Table S1 Loading clusters according to the First/Second components**
A maximum of three codon usages among 54 genera and a genus with the maximum for each Three Codon DNA 9-mer is indicated in the column- three_max and three_genus, respectively.

**Supplementary Table S2 Correlations of a pair of genera based on trimer usage bias**
FS 1
FS 2
FS 3
FS 4
FS 5
FS 6
FS 7
FS 8
FS 9

**Supplementary Table S3 The mean of the correlations in 9 FSs**
The mean and the standard deviation of the correlations of a pair of genera were calculated based on three codon usage (left) and on trimer usage bias (right) in 9 FSs in taxonomic groups- bacterial phyla, AMF, a group of 7 Dikarya, and *Phanerochaete.*

**Supplementary Table S4 The mean and the standard deviation of the correlations of 9 FSs**
Based on three codon usage
Based on trimer usage bias

**Supplementary Table S5 Genus clusters of five additional sequences**

**Supplementary Table S6 Loading clusters of five additional sequences**
