## Supplementary Table S3 for "SeSaMe PS Function: Functional Analysis of the Whole Metagenome Sequencing Data of the Arbuscular Mycorrhizal Fungi"

|  | **Firmicutes** | **Cyanobac** | **Proteobacteria** | **Actinobac** | **AMF** | **7 Dikarya** | **Phanero** | **Mean (std.dev)** |
| --- | --- | --- | --- | --- | --- | --- | --- | --- |
| **Fi** | 0.79:0.93  0.94:0.99  0.90:0.94  0.66:0.17  0.90:0.95  0.59:0.79  0.50:0.77  0.83:0.75  0.96:1.0 | 0.77:0.87  0.90:0.97  0.78:0.80  0.13:-0.14  0.88:0.91  0.61:0.82  0.44:0.78  0.70:0.73  0.94:0.98 | 0.18:0.26  0.73:0.90  0.54^*b^:0.56^*b^  0.49:0.22  0.50:0.40  0.38:0.72  0.16:0.53  0.64:0.56  0.91^*b^:0.96^*b^ | -0.12:-0.17  0.81^*a^:0.93^*a^  0.30^*c^:0.21^*c^  0.10^*d^:-0.097^*d^  0.20^*e^:0.073^*e^  0.11:0.43  -0.13^*f^:-0.12^*f^  0.52^*g^:0.27^*g^  0.51^*h^:0.45^*h^ | 0.69:0.90  0.81:0.94  0.92:0.96  0.82:-0.41  0.77:0.81  0.51:0.75  0.64:0.87  0.90:0.83  0.92:1.0 | 0.38:0.70  0.81:0.95  0.71:0.65  0.50:0.16  0.61:0.50  0.20:0.74  0.42:0.79  0.70:0.56  0.90:0.99 | 0.06:0.065  0.79:0.95  0.34:0.30  -0.24:-0.24  0.44:0.26  -0.18:0.37  -0.19:0.49  0.60:0.36  -0.18:-0.22 | 0.39 (0.37):0.49 (0.47)  0.83 (0.070):0.94 (0.028)  0.64 (0.26):0.63 (0.30)  0.35 (0.37):-0.049 (0.24)  0.61 (0.26):0.56 (0.34)  0.31 (0.29):0.66 (0.18)  0.26 (0.32):0.59 (0.34)  0.70 (0.13):0.58 (0.21)  0.71 (0.42):0.74 (0.47) |
| **Cy** | 0.77:0.87  0.90:0.97  0.78:0.80  0.13:-0.14  0.88:0.91  0.61:0.82  0.44:0.78  0.70:0.73  0.94:0.98 | 0.86:0.93  0.92:0.98  0.93:0.87  0.43:0.16  0.99:0.92  0.81:0.96  0.69:0.88  0.67:0.83  0.97:0.99 | 0.17:0.24  0.66:0.89  0.45^*b^:0.44^*b^  0.29:-0.16  0.54:0.40  0.54:0.85  0.11:0.58  0.52:0.65  0.91^*b^:0.95^*b^ | -0.2:-0.22  0.70^*a^:0.92^*a^  0.16^*c^:0.075^*c^  0.50^*d^:0.14^*d^  0.22^*e^:0.047^*e^  0.25:0.49  -0.22^*f^:-0.15^*f^  0.33^*g^:0.27^*g^  0.56^*h^:0.49^*h^ | 0.61:0.82  0.74:0.92  0.78:0.83  0.14:0.36  0.72:0.75  0.46:0.84  0.69:0.90  0.71:0.85  0.95:0.98 | 0.4:0.71  0.72:0.94  0.41:0.36  0.48:-0.10  0.59:0.45  0.43:0.86  0.55:0.88  0.50:0.53  0.94:0.99 | 0.1:-0.034  0.70:0.94  0.080:0.0048  -0.65:0.065  0.40:0.20  0.028:0.41  -0.017:0.68  0.40:0.26  -0.016:-0.090 | 0.39 (0.38):0.48 (0.47)  0.76 (0.10):0.94 (0.031)  0.51 (0.33):0.48 (0.36)  0.18 (0.40):0.045 (0.19)  0.62 (0.27):0.53 (0.34)  0.45 (0.25):0.75 (0.21)  0.32 (0.36):0.65 (0.37)  0.55 (0.15):0.59 (0.25)  0.75 (0.37):0.76 (0.42) |
| **Pr** | 0.18:0.26  0.73:0.90  0.54^*b^:0.56^*b^  0.49:0.22  0.50:0.40  0.38:0.72  0.16:0.53  0.64:0.56  0.91^*b^:0.96^*b^ | 0.17:0.24  0.66:0.89  0.45^*b^:0.44^*b^  0.27:-0.16  0.54:0.40  0.54:0.85  0.11:0.58  0.52:0.65  0.91^*b^:0.95^*b^ | 0.66:0.57  0.79:0.91  0.34^*b^:0.37^*b^  0.43:0.32  0.70:0.60  0.69:0.83  0.38:0.50  0.80:0.70  0.89^*b^:0.92^*b^ | 0.69:0.62  0.86^*a^:0.95^*a^  0.20^*bc^:0.13^*bc^  0.28^*d^:-0.074^*d^  0.49^*e^:0.42^*e^  0.36:0.47  0.14^*f^:-0.0047^*f^  0.81^*g^:0.48^*g^  0.50^*bh^:0.45^*bh^ | -0.07:0.21  0.68:0.87  0.56^*b^:0.58^*b^  0.59:-0.53  0.37:0.36  0.27:0.73  0.088:0.61  0.70:0.69  0.88^*b^:0.96^*b^ | 0.75:0.57  0.87:0.95  0.46^*b^:0.51^*b^  0.53:0.24  0.68:0.46  0.67:0.85  0.37:0.65  0.87:0.70  0.89^*b^:0.95^*b^ | 0.79:0.68  0.87:0.95  0.28^*b^:0.35^*b^  -0.43:-0.46  0.73:0.58  0.16:0.39  0.36:0.51  0.85:0.53  -0.12^*b^:-0.18^*b^ | 0.45 (0.35):0.45 (0.20)  0.78 (0.090):0.92 (0.033)  0.41 (0.13):0.41 (0.16)  0.31 (0.35):-0.063 (0.34)  0.57 (0.13):0.46 (0.093)  0.44 (0.20):0.69 (0.19)  0.23 (0.13):0.48 (0.22)  0.74 (0.13):0.62 (0.092)  0.69 (0.39):0.71 (0.44) |
| **Ac** | -0.12:-0.17  0.81^*a^:0.93^*a^  0.30^*c^:0.21^*c^  0.10^*d^:-0.097^*d^  0.20^*e^:0.073^*e^  0.11:0.43  -0.13^*f^:-0.12^*f^  0.52^*g^:0.27^*g^  0.51^*h^:0.45^*h^ | -0.2:-0.22  0.70^*a^:0.92^*a^  0.16^*c^:0.075^*c^  0.50^*d^:0.14^*d^  0.22^*e^:0.047^*e^  0.25:0.49  -0.22^*f^:-0.15^*f^  0.33^*g^:0.27^*g^  0.56^*h^:0.49^*h^ | 0.69:0.62  0.86^*a^:0.95^*a^  0.20^*bc^:0.12^*bc^  0.28^*d^:-0.074^*d^  0.49^*e^:0.42^*e^  0.36:0.47  0.14^*f^:-0.0047^*f^  0.81^*g^:0.48^*g^  0.50^*bh^:0.45*^bh^ | 1:0.99  0.99^*a^:1.0^*a^  0.51^*c^:0.31^*c^  0.65^*d^:0.29^*d^  0.49^*e^:0.49^*e^  0.28:0.31  0.44^*f^:0.36^*f^  0.97^*g^:0.78^*g^  0.50^*h^:0.48^*h^ | -0.26:-0.19  0.75^*a^:90^*a^  0.25^*c^:0.20^*c^  0.098^*d^:0.26^*d^  0.077^*e^:0.16^*e^  0.17:0.50  -0.20^*f^:-0.15^*f^  0.61^*g^:0.44^*g^  0.60^*h^:0.47^*h^ | 0.66:0.33  0.97^*a^:1.0^*a^  0.16^*c^:0.20^*c^  0.53^*d^:-0.035^*d^ 0.42^*e^:0.37^*e^  0.38:0.51  -0.11^*f^:-0.12^*f^  0.94^*g^:0.71^*g^  0.52^*h^:0.47^*h^ | 0.85:0.92  0.96^*a^:1.0^*a^  -0.034^*c^:0.15^*c^  -0.76^*d^:-0.12^*d^  0.54^*e^:0.57^*e^  0.049:0.19  0.33^*f^:0.098^*f^  0.96^*g^:0.66^*g^  0.094^*h^:0.1^5*h^ | 0.37 (0.54):0.33 (0.53)  0.86 (0.12):0.96 (0.042)  0.22 (0.17):0.18 (0.073)  0.20 (0.48):0.052 (0.17)  0.35 (0.18):0.30 (0.21)  0.23 (0.12):0.41 (0.12)  0.035 (0.27):-0.012 (0.19)  0.74 (0.25):0.51 (0.21)  0.47 (0.17):0.42 (0.12) |
| **AMF** | 0.69:0.90  0.81:0.94  0.93:0.96  0.82:-0.41  0.77:0.81  0.51:0.75  0.64:0.87  0.90:0.83  0.92:1.0 | 0.61:0.82  0.74:0.92  0.78:0.83  0.14:0.36  0.72:0.75  0.46:0.84  0.69:0.90  0.71:0.85  0.95:0.99 | -0.07:0.21  0.68:0.87  0.56^*b^:0.58^*b^  0.59:-0.53  0.37:0.36  0.27:0.73  0.088:0.62  0.70:0.69  0.88^*b^:0.96^*b^ | -0.26:-0.19  0.75^*a^:90^*a^  0.25^*c^:0.20^*c^  0.098^*d^:0.26^*d^  0.077^*e^:0.16^*e^  0.17:0.50  -0.20^*f^:-0.15^*f^  0.61^*g^:0.44^*g^  0.60^*h^:0.47^*h^ | 1.0:1.0  1.0:1.0  1.0:1.0  1.0:1.0  1.0:1.0  1.0:1.0  1.0:1.0  1.0:1.0  1.0:1.0 | 0.09:0.65  0.79:0.92  0.84:0.67  0.59:-0.36  0.55:0.76  -0.023:0.76  0.57:0.92  0.79:0.74  0.92:0.98 | -0.25:-0.091  0.70:0.90  0.52:0.33  -0.25:0.53  0.46:0.57  -0.73:0.12  -0.32:0.56  0.70:0.51  0.11:-0.20 | 0.26 (0.50):0.47 (0.49)  0.78 (0.11):0.92 (0.041)  0.70 (0.26):0.65 (0.30)  0.43 (0.44):0.12 (0.57)  0.56 (0.30):0.63 (0.29)  0.23 (0.54):0.67 (0.29)  0.35 (0.50):0.67 (0.40)  0.77 (0.13):0.72 (0.20)  0.77 (0.32):0.74 (0.46) |
| **7 Di** | 0.38:0.70  0.81:0.95  0.71:0.65  0.50:0.16  0.61:0.50  0.20:0.74  0.42:0.79  0.70:0.56  0.90:0.99 | 0.4:0.71  0.72:0.94  0.41:0.36  0.48:-0.10  0.59:0.45  0.43:0.86  0.55:0.88  0.50:0.53  0.94:0.99 | 0.75:0.58  0.87:0.95  0.46^*b^:0.51^*b^  0.53:0.24  0.68:0.46  0.67:0.85  0.37:0.65  0.87:0.70  0.89^*b^:0.95^*b^ | 0.66:0.33  0.97^*a^:1.0^*a^  0.16^*c^:0.20^*c^  0.53^*d^:-0.035^*d^  0.42^*e^:0.37^*e^  0.38:0.51  -0.11^*f^:-0.12^*f^:  0.94^*g^:0.71^*g^  0.52^*h^:0.47^*h^ | 0.09:0.65  0.79:0.92  0.84:0.67  0.59:-0.37  0.55:0.76  -0.023:0.76  0.57:0.92  0.79:0.74  0.92:0.99 | 0.91:0.88  0.99:1.0  0.94:0.96  0.73:0.18  0.90:0.92  0.84:0.95  0.78:0.97  0.99:0.98  0.94:0.99 | 0.88:0.52  0.98:1.0  0.80:0.88  -0.75:-0.36  0.84:0.80  0.46:0.54  0.24:0.75  0.98:0.92  0.061:-0.13 | 0.58 (0.30):0.62 (0.17)  0.87 (0.11):0.97 (0.030)  0.62 (0.28):0.60 (0.27)  0.37 (0.50):-0.043 (0.25)  0.66 (0.17):0.61 (0.21)  0.42 (0.29):0.75 (0.17)  0.40 (0.28):0.69 (0.37)  0.82 (0.17):0.74 (0.17)  0.74 (0.34):0.75 (0.43) |
| **Ph** | 0.06:-0.065  0.79:0.95  0.34:0.30  -0.23:-0.24  0.44:0.26  -0.18:0.37  -0.19:0.48  0.60:0.36  -0.18:-0.22 | 0.1:-0.034  0.70:0.94  0.080:0.0048  -0.65:0.065  0.40:0.20  0.028:0.41  -0.017:0.68  0.40:0.26  -0.016:-0.090 | 0.79:0.68  0.87:0.95  0.28^*b^:0.35^*b^  -0.43:-0.46  0.73:0.58  0.16:0.39  0.35:0.51  0.85:0.53  -0.12^*b^:-0.18^*b^ | 0.85:0.92  0.96^*a^:1.0^*a^  -0.034^*c^:0.15^*c^  -0.76^*d^:-0.12^*d^  0.54^*e^:0.57^*e^  0.049:0.19  0.33^*f^:0.098^*f^:  0.96^*g^:0.66^*g^  0.094^*h^:0.15^*h^ | -0.25:-0.091  0.70:0.90  0.52:0.33  -0.25:0.53  0.46:0.57  -0.73:0.11  -0.32:0.56  0.70:0.51  0.11:-0.20 | 0.88:0.52  0.98:1.0  0.80:0.88  -0.75:-0.36  0.84:0.80  0.46:0.54  0.24:0.75  0.98:0.93  0.061:-0.13 | 1.0:1.0  1.0:1.0  1.0:1.0  1.0:1.0  1.0:1.0  1.0:1.0  1.0:1.0  1.0:1.0  1.0:1.0 | 0.49 (0.50):0.42 (0.48)  0.86 (0.13):0.96 (0.037)  0.43 (0.37):0.43 (0.37)  -0.30 (0.61):0.060 (0.52)  0.62 (0.23):0.57 (0.28)  0.11 (0.54):0.43 (0.29)  0.20 (0.44):0.58 (0.28)  0.78 (0.23):0.61 (0.28)  0.14 (0.40):0.047 (0.44) |

**Supplementary Table 3 The mean of the correlations in 9 FSs**

The mean and the standard deviation of the correlations of a pair of genera were calculated based on three codon usage (left) and on trimer usage bias (right) in 9 FSs in taxonomic groups- bacterial phyla, AMF, a group of 7 Dikarya, and *Phanerochaete.*

*Note*: Red color indicates the mean of the correlations of a pair of genera belonging to the same taxonomic group. Brown color indicates the mean and the standard deviation of 7 taxonomic groups per FS. Abbreviations: Fi:Firmicutes; Cy,Cyanobac:Cyanobacteria; Pr:Proteobacteria; Ac,Actinobac:Actinobacteria; Ph,Phanero:Phanerochaete. * indicates that the correlation between the indicated genus below and each of 54 genera was NaN (Not A Number) due to having zero value. *a: Kocuria, Microbacterium, Pseudonocardia; *b: Azorhizobium; *c: Microbacterium, Micrococcus, Pseudonocardia; *d: Kocuria, Micrococcus, Pseudonocardia; *e: Microbacterium, Micrococcus; *f: Micrococcus, Pseudonocardia; *g: Kocuria, Microbacterium; *h: Kocuria, Microbacterium, Micrococcus
