## Supplementary Table S5 for "SeSaMe PS Function: Functional Analysis of the Whole Metagenome Sequencing Data of the Arbuscular Mycorrhizal Fungi"

|  | **seq252** | **seq284** | **seq337** | **seq475** | **seq528** |
| --- | --- | --- | --- | --- | --- |
| **Cluster 0** | 0,1,2,4,5,9,10,16,17,19,21,22,23,24,25,26,29,30,34,35,36,37,40,41,42,43,44,49,51 | 4,5,17,22,34,37,41,43,44 | 1,4,5,9,10,16,19,21,22,23,24,25,26,30,34,35,36,37,40,41,42,43,44,51 | 7,13,28,33 | 1,2,4,5,8,9,10,16,19,21,22,23,24,25,26,30,34,35,36,37,40,41,42,43,44,51 |
| **Cluster 1** | 32,39 | 3,12,20,31,38 | 12,20 | 11,45 | 11,45 |
| **Cluster 2** | 38 | 7,13,27,32,39,47,50,52,53 | 13,31 | 4,5,9,10,16,19,22,23,24,25,30,34,35,37,41,42,43,44 | 6,12 |
| **Cluster 3** | 6,20 | 11,45 | 38,45 | 27,29,39,47,48,50,52,53 | 3,20 |
| **Cluster 4** | 7,8,14,15,18,46,47,48,52,53 | 6,33 | 27,33 | 6,12 | 27,33 |
| **Cluster 5** | 11,45 | 9 | 7,8,14,15,29,39,47,50,52,53 | 0,1,2,18,21,26,36,40 | 28,32,39 |
| **Cluster 6** | 33,50 | 1,2,21,26,36,40 | 0,2,17,18,46,48,49 | 3,20 | 38 |
| **Cluster 7** | 12 | 28,29 | 3,6 | 38 | 7,13,31 |
| **Cluster 8** | 3,13 | 0,8,14,15,18,46,48,49,51 | 11 | 8,14,15,17,46,49,51 | 14,15,18,29,47,48,50,52,53 |
| **Cluster 9** | 27,28,31 | 10,16,19,23,24,25,30,35,42 | 28,32 | 31,32 | 0,17,46,49 |

**Supplementary Table 5 Genus clusters of five additional sequences**

*Note*:0:Acidithiobacillus,1:Acidobacterium,2:Agrobacterium,3:Anabaena,4:Azorhizobium,5:Azotobacter,6:Bacillus,7:Bdellovibrio,8:Beijerinckia,9:Bradyrhizobium,10:Caulobacter,11:Clostridium,12:Cyanobacterium,13:Desulfotomaculum,14:Desulfovibrio,15:Erwinia,16:Frankia,17:Geobacter,18:Klebsiella,19:Kocuria,20:Leuconostoc,21:Mesorhizobium,22:Methylococcus,23:Microbacterium,24:Micrococcus,25:Myxococcus,26:Nitrobacter,27:Nitrosococcus,28:Nitrosomonas,29:Nitrosospira,30:Nocardia,31:Nostoc,32:Oscillatoria,33:Pseudanabaena,34:Pseudomonas,35:Pseudonocardia,36:Rhizobium,37:Rhodobacter,38:Rickettsia,39:Shewanella,40:Sinorhizobium,41:Sphingomonas,42:Streptomyces,43:Variovorax,44:Xanthomonas,45:AMF,46:Aspergillus,47:Cenococcum,48:Cryptococcus,49:Mycosphaerella,50:Oidiodendron,51:Phanerochaete,52:Scleroderma,53:Sebacina
