## Supplementary Table S6 for "SeSaMe PS Function: Functional Analysis of the Whole Metagenome Sequencing Data of the Arbuscular Mycorrhizal Fungi"

| **seq252** | **seq284** | **seq337** | **seq475** | **seq528** |
| --- | --- | --- | --- | --- |
| Cluster 0  69_JEJ_VNL_GTGAATTTG,  Cluster: 0 done  Cluster 1  59_EKA_NWK_AATTGGAAA,  Cluster: 1 done  Cluster 2  34_AJE_RAN_AGAGCAAAT,  Cluster: 2 done  Cluster 3  30_GAJ_GKV_GGTAAAGTG,  Cluster: 3 done  Cluster 4  Major cluster  Cluster: 4 done  Cluster 5  37_KKC_WFD_TGGTTTGAT,  Cluster: 5 done  Cluster 6  1_BDK_HTF_CATACATTC,  Cluster: 6 done  Cluster 7  70_EJA_NLK_AATTTGAAA,  71_JAD_LKS_TTGAAAAGT,  Cluster: 7 done  Cluster 8  26_ACC_KDE_AAAGATGAG,  Cluster: 8 done  Cluster 9  18_JKC_LYD_TTGTATGAC,  Cluster: 9 done  Cluster 10  31_AJK_KVY_AAAGTGTAT,  Cluster: 10 done  Cluster 11  40_GEJ_GNI_GGGAACATA,  Cluster: 11 done  Cluster 12  36_EKK_NWF_AATTGGTTT,  Cluster: 12 done | Cluster 0  74_HGA_PGK_CCAGGAAAG,  Cluster: 0 done  Cluster 1  36_JJD_IVT_ATAGTGACT,  Cluster: 1 done  Cluster 2  27_JHJ_LPI_TTGCCGATT,  Cluster: 2 done  Cluster 3  52_CHJ_EPI_GAACCAATC,  64_JJD_ILS_ATTTTGTCA,  Cluster: 3 done  Cluster 4  1_GEH_GNP_GGTAACCCA,  Cluster: 4 done  Cluster 5  Major cluster  Cluster: 5 done  Cluster 6  43_AJA_KLR_AAACTACGC,  Cluster: 6 done  Cluster 7 26_CJH_ELP_GAATTGCCG,  Cluster: 7 done  Cluster 8 17_AAE_RRN_CGTAGAAAC,  Cluster: 8 done  Cluster 9 37_JDC_VTE_GTGACTGAA,  Cluster: 9 done  Cluster 10 45_AKA_RYR_CGCTATAGA,  Cluster: 10 done  Cluster 11 57_KIE_YMQ_TATATGCAA,  Cluster: 11 done  Cluster 12 78_KJI_YLM_TATCTCATG,  Cluster: 12 done | Cluster 0  12_KKJ_YFL_TATTTTCTC,  Cluster: 0 done  Cluster 1  Major cluster  Cluster: 1 done  Cluster 2  8_DDK_TSF_ACATCTTTT,  Cluster: 2 done  Cluster 3  46_ICJ_MDA_ATGGATGCA,  Cluster: 3 done  Cluster 4  40_AKC_KFE_AAATTTGAG,  Cluster: 4 done  Cluster 5  52_AJE_KLQ_AAACTTCAA,  Cluster: 5 done  Cluster 6  27_AEG_RNG_CGTAATGGG,  Cluster: 6 done  Cluster 7  2_JJD_LVS_TTGGTATCA,  Cluster: 7 done  Cluster 8  5_DAK_TKY_ACTAAGTAT,  Cluster: 8 done  Cluster 9  22_EKJ_QFI_CAATTCATA,  Cluster: 9 done  Cluster 10  4_DDA_STK_TCAACTAAG,  Cluster: 10 done  Cluster 11  53_JEJ_LQI_CTTCAAATA,  Cluster: 11 done  Cluster 12  7_KDD_YTS_TATACATCT,  35_KCH_YEP_TATGAACCA,  36_CHJ_EPI_GAACCAATT,  37_HJA_PIK_CCAATTAAA,  Cluster: 12 done | Cluster 0  51_DDD_SSS_AGTTCGAGT,  Cluster: 0 done  Cluster 1  45_DGD_SGT_TCAGGTACT,  Cluster: 1 done  Cluster 2  43_GDD_GTS_GGTACTTCA,  Cluster: 2 done  Cluster 3  24_EAD_NKS_AATAAATCA,  56_EAK_QKY_CAAAAATAT,  Cluster: 3 done  Cluster 4  11_CAJ_DKL_GACAAATTA,  26_DJC_SAE_TCAGCTGAA,  Cluster: 4 done  Cluster 5  Major cluster  Cluster: 5 done  Cluster 6  40_AJA_RLR_AGATTACGA,  Cluster: 6 done  Cluster 7  27_JCE_AEQ_GCTGAACAG,  Cluster: 7 done  Cluster 8  28_CEG_EQG_GAACAGGGA,  Cluster: 8 done  Cluster 9  14_CCD_DDT_GACGACACA,  Cluster: 9 done  Cluster 10  33_CJA_EAR_GAAGCTAGG,  Cluster: 10 done  Cluster 11  29_EGE_QGN_CAGGGAAAT,  Cluster: 11 done  Cluster 12  37_DKK_SFY_AGTTTTTAT,  Cluster: 12 done | Cluster 0  7_AGJ_KGA_AAAGGAGCA,  Cluster: 0 done  Cluster 1  Major cluster  Cluster: 1 done  Cluster 2  24_AEJ_KNA_AAAAATGCG,  Cluster: 2 done  Cluster 3  31_JDA_ASK_GCCTCGAAA,  Cluster: 3 done  Cluster 4  17_AEK_RNF_AGAAATTTT,  Cluster: 4 done  Cluster 5  25_EJA_NAK_AATGCGAAA,  Cluster: 5 done  Cluster 6  20_EKK_NYY_AATTATTAT,  Cluster: 6 done  Cluster 7  36_AJH_KAP_AAAGCTCCA,  Cluster: 7 done  Cluster 8  30_CJD_DAS_GATGCCTCG,  Cluster: 8 done  Cluster 9  1_CDJ_ESL_GAATCTCTT,  Cluster: 9 done  Cluster 10  11_JJJ_LLI_TTATTAATC,  34_JDA_ITK_ATTACAAAA,  Cluster: 10 done  Cluster 11  5_JDA_ATK_GCTACAAAA,  18_EKE_NFN_AATTTTAAT,  Cluster: 11 done  Cluster 12  13_JED_INT_ATCAACACT,  Cluster: 12 done |

**Supplementary Table 6 Loading clusters of five additional sequences**

*Note*:

seq252:GCACATACATTCTATGAAGTAAATAATGCATTAGAATGGATACCTTATGATAAATTGTATGACATTAAATATATTACGAAAGATGAGTTAGGTAAAGTGTATAGAGCAAATTGGTTTGATGGGAACATAATTGATAAATATTATAGTTATAATTATTGGGGTGATGTATTAAAACATAATTGGAAAAGAAACTATCCTAATATGTTTGTGAATTTGAAAAGTTTAAATTCTCCAAATGATCTTAC ;

seq284:CCAATGGTAACCCAAATGGAAATGATAATGGTAATGGCAATGGTACAGAACGACGTAGAAACGTAGAAGATCTTTATTCTGAATTGCCGATTGATAGTAAAACTAAGGAAATAGTGACTGAAGTTAATGCAAAACTACGCTATAGATATGTAAATATGGAACCAATCAAGCTTTATATGCAAGTTTGCCAATTTATTTTGTCATTATTTCCTGATGTACCGGATCCAGGAAAGTTATATCTCATGTTTCCGGATGGTAAAA ;

seq337:TATTTATTTGGTATCAACTAAGTATACATCTTTTTTATATTTTCTCTTTCCAAAATTAACAAATTTACAATTCATAAGAATACGTAATGGGGATAATATTAATAATTATGAACCAATTAAAAAATTTGAGGAATACGCAATGGATGCAAGTTATTATAAACTTCAAATACTTGAGT ;

seq475:AGCTCAATACAATCTTGGAGTTATTTATGAAACTGACAAATTAGACGACACAATTGCAGCACTGTATTGGTATAATAAATCAGCTGAACAGGGAAATCATGAAGCTAGGGAAAGTTTTTATAGATTACGAGGTACTTCAGGTACTAAGACTGTTAGTTCGAGTAGTATACAAAAATATGGTTCTATGGGTAT ;

seq528:CTTGAATCTCTTCTTGCTACAAAAGGAGCAGAGTTATTAATCAACACTTTAAGAAATTTTAATTATTATAAAAAAAATGCGAAAGAACAAGATGCCTCGAAAATTACAAAAGCTCCAAAAATTAAAAAAGAAATGAGTAAAATTAAGTGGTCACAAATT
